# Glassy phase in dynamically-balanced neuronal networks

**DOI:** 10.1101/2022.03.14.484348

**Authors:** Kevin Berlemont, Gianluigi Mongillo

**Affiliations:** Center for Neural Science, New York University, New York, NY 10003, USA; Sorbonne Université, INSERM, CNRS, Institut de la Vision, F-75012 Paris, France and Centre National de la Recherche Scientifique (CNRS), Paris, France

## Abstract

We present a novel mean-field theory for balanced neuronal networks with arbitrary levels of symmetry in the synaptic connectivity. The theory determines the fixed point of the network dynamics and the conditions for its stability. The fixed point becomes unstable by increasing the synaptic gain beyond a critical value that depends on the level of symmetry. Beyond this critical gain, for positive levels of symmetry, we find a previously unreported phase. In this phase, the dynamical landscape is dominated by a large number of marginally-stable fixed points. As a result, the network dynamics exhibit non-exponential relaxation and ergodicity is broken. We discuss the relevance of such a *glassy* phase for understanding dynamical and computational aspects of cortical operation.

---

Balanced networks are a key theoretical construct to describe the dynamical regime in which the cortex operates and to understand its computational implications [1]. In these networks, neuronal activity dynamically adjusts so that the average input to a neuron is of the same order as the spatial and/or temporal fluctuations [2–5]. This results in a dynamical state – the balanced regime – characterized by low and heterogeneous single-neuron firing rates, with temporally-irregular spiking [6–9] and vanishingly small pairwise correlations in the activity of different neurons [5, 10], consistent with experimental observations in the cortex.

The balanced regime is understood in random networks of binary neurons [3, 5, 10] and of rate-based neurons [11, 12], *random* meaning that there are no correlation in the connectivity matrix. However, local connectivity (i.e., on spatial scales of the order of 100 μm) in the cortex is not random. Reciprocal synaptic connections are both over-represented, as compared to random networks, and stronger than average [13–15]. The effects of such partial symmetry on the dynamics are not well understood [16–18], because the analytical description is difficult. In fact, fluctuations in the synaptic inputs are not Gaussian, due to the correlations in the connectivity. Currently, there is no mean-field description for partially-symmetric balanced networks.

In this letter, we show that the non-Gaussianity of the inputs can be conveniently dealt with by a *cavity* approximation [19], which leads to an exact mean-field theory when correlations are restricted to reciprocal connections. The theory determines the distribution of neuronal activity in the fixed point and its stability. When the mean-field solution is unstable, we study numerically the network dynamics. We find that the dynamics are not ergodic, provided the correlation in the connectivity is positive.

We consider a fully connected network of *N* inhibitory rate-based neurons. The state of neuron *i* is described by its synaptic input, *h_i_*, which evolves in time (in units of the time constant) according to

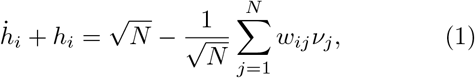

where the first term on the r.h.s. describes an excitatory external input, *w_ij_* > 0 is the efficacy of the synapse connecting neuron *j* to neuron *i*, and *ν_j_*, the rate of neuron *j*, is a function of its synaptic input, i.e., *ν_j_* = *ϕ*(*h_j_*). We choose *ϕ*(*h_j_*) ≡ *h_j_*Θ(*h_j_*), where Θ(·) is the Heaviside function. The *w_ij_* are given by

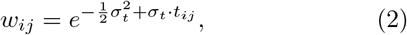

where *t_ij_* is normally distributed with mean zero and unit variance. Thus, the synaptic efficacies are log-normally distributed with mean 〈*w*〉 = 1 and variance (synaptic gain) 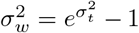, which is tuned by adjusting 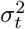. The correlation of reciprocal connections (level of symmetry), *ρ_w_*, is tuned by adjusting the correlation *ρ_t_* = 〈*t_ij_t_ji_*).

The model includes elements of biological realism lacking in previous studies [16, 17]: (i) the synaptic efficacies are sign-constrained, to comply with the so-called Dale’s law [20]; (ii) the single-neuron activity is described by a non-negative variable. The network, then, operates in the balanced regime for *N* → ∞. The other modeling choices, such as the distribution of the *w_ij_* and the specific *ϕ*(·), have been motivated by considerations of analytical/numerical tractability of the resulting equations. They are immaterial for the theory developed below.

We are interested in a mean-field description of the fixed point(s) of our model network (see [21] for a detailed derivation). For this, we need to determine the distribution of the synaptic inputs in a fixed point. We cannot use the central limit theorem to evaluate the sum in Eq. (1) because all *ν_j_* for *j* ≠ *i* depend on *ν_i_* and, hence, they are not independent. Moreover, when *ρ_w_* ≠ 0, *w_ij_* and *ν_j_* are also correlated because *ν_j_* depends on *w_ji_ν_i_*, and *w_ij_* and *w_ji_* are correlated. To make progress, we assume that the *ν_j_* are correlated *only* through *ν_i_*, and rewrite *h_i_* as

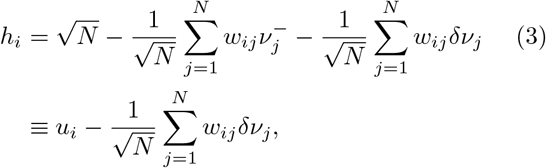

where 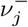 is the rate of neuron *j* when the outgoing connections from neuron *i* are removed (i.e., *w_ji_* = 0 for all *j*) and, hence, *δν_j_* is the change in the rate of neuron *j* due to neuron *i*. By construction the 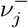 are now independent and the distribution of the *cavity input, u_i_*, across realizations of the *w_ij_* becomes Gaussian in the limit *N* → ∞. The corresponding mean, *μ*, and variance, *σ*^2^, can be expressed in terms of the means and variances of the *w_ij_* and the 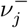. This is because the *w_ij_* and the 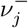 are also independent by construction.

The second sum on the r.h.s. of Eq. (3) – the *reaction term* – can be computed with linear response theory, because the presence of neuron *i* induces only a small (i.e., 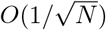) perturbation in the input to neuron *j*. In the limit *N* → ∞, we find

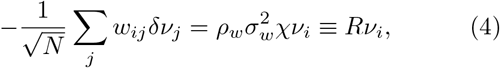

with *χ* ≡ *N*^−1^∑_*j*_ *χ_jj_*, where *χ_jj_* is the static *local* susceptibility of neuron *j*. Finally, we can determine self-consistently *h_i_* using Eqs. (3) and (4), i.e.,

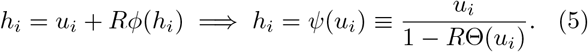

As anticipated, the distribution of the synaptic inputs in the fixed point is non-Gaussian when *ρ_w_* ≠ 0. However, Eq. (5) solves the problem. We can express *ν_i_* as a function of the cavity input *u_i_*, i.e., 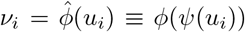. Using the effective transduction function, 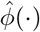, we then determine the mean and the variance of the distribution of the cavity inputs, as in a random network, by requiring self-consistency. Additionally, however, we impose self-consistency on *χ*, which determines in turn the effective transduction function self-consistently.

The mean-field theory is valid only if the fixed point is asymptotically stable, otherwise we cannot use linear response theory to compute the reaction term. The stability of the fixed point can be assessed by computing the leading order corrections to the mean-field theory, i.e., the *δν_j_* in Eq. (3). If these adjustments are *non-physical*, the mean-field solution, and hence the associated fixed point, is unstable [19]. Using 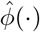 we can write 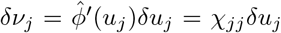 (to leading order), where *δu_j_* is the change in the cavity input to neuron *j* due to neuron *i*. The key simplification is that the *δu_j_* are Gaussian (by construction). Determining self-consistently the *δν_j_* reduces to a standard problem [21]. We find

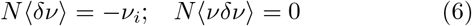

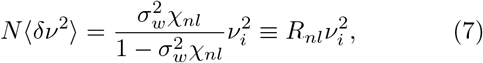

where we have defined 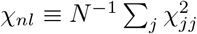. Clearly, 〈*δν*^2^〉 must be non-negative and finite, which requires 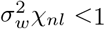. This latter condition is satisfied whenever

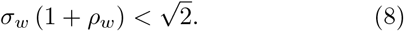

As long as the mean-field solution is stable, there is a unique, stable fixed point of Eq. (1) [21].

Next, we compare the theoretical predictions with the results of numerical simulations. We find a very good agreement as illustrated in Fig. 1 for the mean-field theory, and in Fig. 2 for the leading order corrections to the mean-field theory, i.e., Eqs. (6)–(7). As can be seen in Fig. 1d, the fraction of active neurons (i.e., with *ν_i_* > 0) in the fixed point, *f*, decreases with increasing *ρ_w_*. This can be understood as follows. The stability of the fixed point is controlled by the spectrum of the Jacobian, 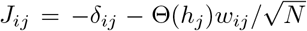, where *δ_ij_* is the Kronecker symbol. The average spectrum (i.e., over realizations of the synaptic matrix) is easily evaluated in the limit *N* → ∞. There is one real, negative eigenvalue 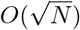, (1 − *f*)(*N* − 1) (on average) eigenvalues equal to −1, and the remaining *f*(*N* − 1) eigenvalues, *λ_k_* – which determine the stability – are *O*(1) and uniformly distributed within an ellipse in the complex plane [22]. Their real part satisfies

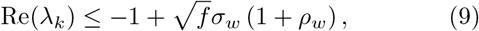

for all *k* with probability 1 when *N* → ∞. Thus, decreasing f has a stabilizing effect that counteracts the destabilizing effect of increasing *ρ_w_*. We can use Eq. (9) to upper bound f in a stable fixed point, for given *ρ_w_* and 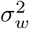. It must be *f* ≤ *f*\*, where

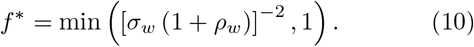

**Figure 1.**
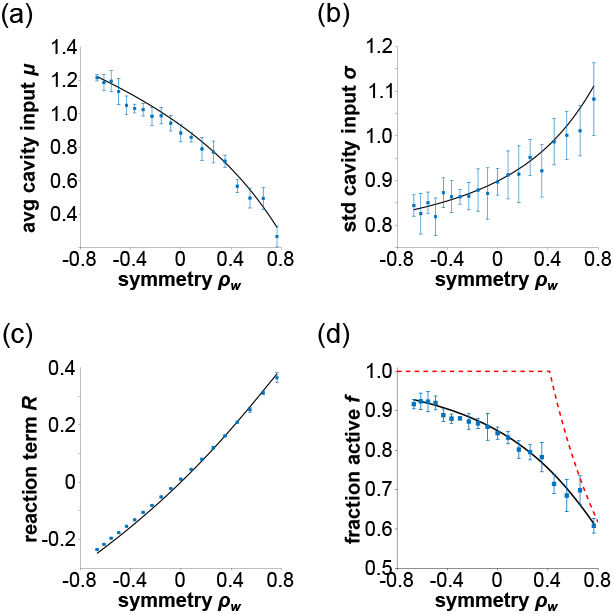
Comparison between the predictions of the mean-field theory (black, full lines) and the results of the numerical simulations (blue symbols, with error bars) at varying *ρ_w_*, for 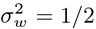. Error bars represent 95% confidence intervals. The red, dashed line in (d) is the stability bound *f*\* in Eq. 10. Network size: *N* = 4000.

**Figure 2.**
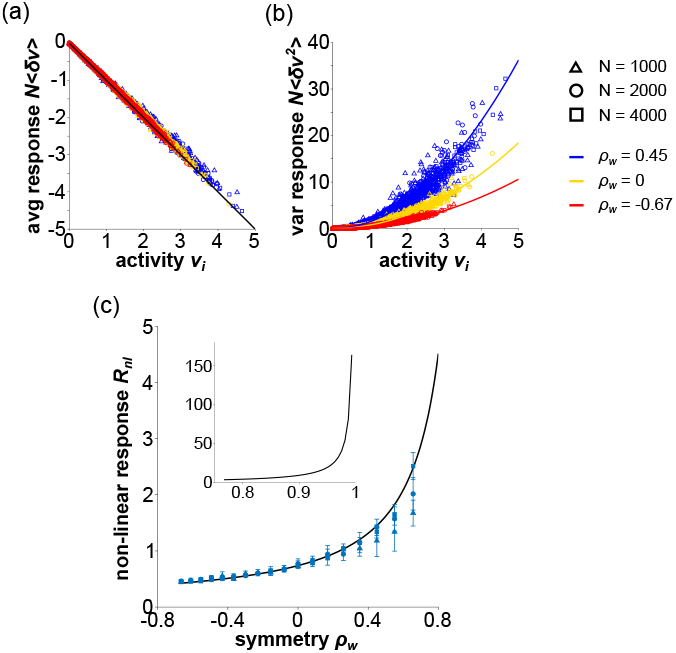
Comparison between the theoretical corrections to the mean-field theory (full lines) and the results of numerical simulations (symbols) at varying *ρ_w_*, for 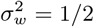. Error bars represent 95% confidence intervals. The inset in (c) shows the divergence of the non-linear response for *ρ_w_* → 1.

In Fig. 1d we plot *f*\* as a function of *ρ_w_* (red, dashed line). It is always *f* < *f*\*, and *f* = *f*\* = 1/2 at the critical point.

What happens when the mean-field solution is unstable? For *ρ_w_* = 1, the dynamics are governed by an *energy* function featuring exponentially many minima (see, e.g., [23]) and the network is multi-stable. By contrast, for *ρ_w_* = 0 there are no stable fixed points and the dynamics are chaotic [11, 12, 24–26]. Hence, between *ρ_w_* = 1 and *ρ_w_* = 0 there is a transition between an exponential number of stable fixed points and no stable fixed point. We try to locate the transition by numerically integrating Eq. (1) for 2000 time constants while starting from 100 different initial conditions (ICs), chosen randomly and independently. This procedure is repeated for 5 realizations of the connectivity matrix at given *ρ_w_* and 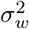.

We start with small networks, i.e., *N* ≤ 800. The rationale for this choice should be clear shortly. The results are illustrated in Fig. (3). In Fig. 3a, we plot the number of fixed points found as a function of the *rescaled* synaptic gain, 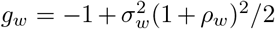, for three values of *ρ_w_* > 0. As predicted by the theory, different ICs lead to the same fixed point for *g_w_* < 0, i.e., when the mean-field solution is stable (see Eq. (8)). Instead, when *g_w_* > 0 (i.e., the mean-field solution is unstable), the network is multi-stable for *ρ_w_* ≥ 0.5. For *ρ_w_* < 0.5 (and *g_w_* > 0), the dynamics rarely converged to a fixed point within the allotted time (data not shown). Obviously, this does not locate the transition. There could be an exponential number of fixed points for *N* → ∞ and yet none at a given finite *N*, depending on how fast the logarithm of the *typical* number of fixed points grows with *N*.

**Figure 3.**
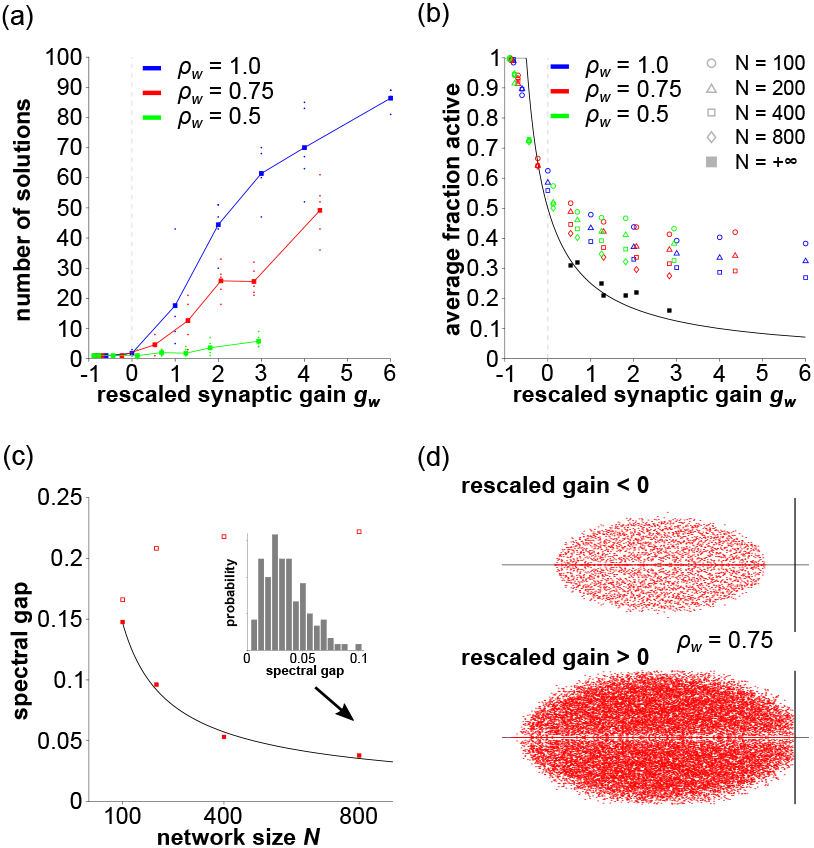
(a) Average number of fixed points and (b) Average fraction of active neurons in a fixed point as a function of the rescaled synaptic gain *g_w_* for three values of *ρ_w_*. Symbols denote network size *N*. The dots in (a) denote the number of fixed points for one realization of the synaptic matrix. The black curve in (b) is the stability bound *f*\* in Eq. 10. (c) Average spectral gap as a function of the network size *N* for *g_w_* = −0.5 (red, open squares) and *g_w_* = 2.0 (red, full squares) at *ρ_w_* = 0.75. The black line is the least-squares fitting to *αN*^−2/3^. The inset shows the variability of the spectral gap across the different fixed points found for *g_w_* = 2, *ρ_w_* = 0.75 and *N* = 800. (d) Sample spectra of the Jacobian in the fixed point(s) for *g_w_* = −0.5 (top) and *g_w_* = 2 (bottom). In both cases, *ρ_w_* = 0.75 and *N* = 800.

In Fig. 3b, we plot *f* averaged over fixed points and connectivity matrices, 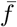, as a function of *g_w_*. Consistently with the stability bound Eq. (10), 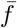 decreases with *g_w_*. However, it is systematically above *f*\* for *g_w_* ≥ 0, which is indicative of strong finite-size effects. In fact, 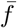 decreases with *N* at parity of *g_w_*. Thus, we performed a finite-size scaling analysis. Interestingly, 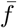 extrapolated at *N* → ∞ is very close to *f*\*, suggesting that the (vast) majority of the fixed points found starting from random ICs are marginally stable for *N* → ∞. This is confirmed by the finite-size scaling analysis of the *spectral gap* – the distance of the spectrum of the Jacobian from the imaginary axis – illustrated in Fig. 3c. There, we plot the spectral gap averaged over fixed points as a function *N*, for *ρ_w_* = 0.75 and two values of *g_w_*. For *g_w_* = −0.5, the average spectral gap rapidly converges to a positive value. By contrast, for *g_w_* = 2 the average spectral gap keeps approaching 0. This decay is well fitted by *αN*^−2/3^ (*α* is a free parameter), as expected if the spectrum *touches* the imaginary axis for *N* → ∞ [27]. The inset illustrates the variability of the spectral gap across fixed points, confirming that, indeed, most fixed points are at the edge of stability. For purpose of illustration, we plot in Fig. 3d the spectrum of the Jacobian for the fixed point found at *g_w_* = −0.5 (top) and for the fixed points found at *g_w_* = 2 (bottom). Note that *g_w_* = 2 is far from the critical line.

These results are explained naturally if, for *ρ_w_* > 0, the number of stable fixed points of Eq. (1) grows exponentially with *N*. At a given *ρ_w_*, the marginally-stable fixed points are exponentially more than the asymptotically-stable ones. This explains why one always finds close-to-marginal fixed points when starting from random ICs. For the same *ρ_w_*, however, there must be also an exponential number of asymptotically-stable fixed points. This is needed to explain why the marginality of the dynamics is structurally stable against decreasing *ρ_w_*. The number of stable fixed points decreases with decreasing *ρ_w_* until only unstable fixed points are left at *ρ_w_* = 0 (and 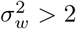). This leads to the chaotic phase [26].

If the above scenario is correct, we expect broken ergodicity for any *ρ_w_* > 0 via the same mechanisms at work in mean-field models of glass forming materials [19, 28, 29]. Note, however, that one can no longer define an energy function to describe the dynamics when *ρ_w_* < 1. Thus, there is no guarantee that the fixed points and their stability *alone* determine the dynamics of the network. Nevertheless, it seems plausible (to us) that if other attractors existed (e.g., limit cycles), they would have vanishing basins of attraction for *N* → ∞ because, in the same limit, an exponential number of stable fixed points has to be accommodated in the phase space. The results we present now support this conjecture.

By definition, the network dynamics are ergodic if the average of the *h_i_* over a suitably long time interval are independent of the initial conditions. We define

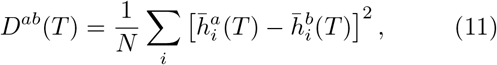

where 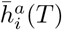 is the time-average of *h_i_* over a time interval *T* when the dynamics Eq. (1) start from the IC *a* (specifying 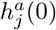 for all *j*). Thus, the dynamics are ergodic if lim_*T*→∞_ *D^ab^*(*T*) = 0 for all pairs (*ab*) of different ICs. We numerically estimated the behavior of *D^ab^*(*T*). The results are shown in Fig. 4. When *g_w_* < 0, *D^ab^*(*T*) rapidly goes to 0, regardless of *ρ_w_*. When *g_w_* > 0, *D^ab^*(*T*) goes to 0 for *ρ_w_* ≤ 0 (Fig. 4b), and to a finite value for *ρ_w_* = 1 (Fig. 4c). Interestingly, for 0 < *ρ_w_* < 1, the decay of *D^ab^*(*T*) is so slow that *D^ab^*(2000) > 0 for both *ρ_w_* = 0.26 (Fig. 4(d)) and *ρ_w_* = 0.45 (Fig. 4(e)). We have fitted *D^ab^*(*T*), averaged over (*ab*), to a stretched exponential with an offset. As can be seen, the stretched exponential provides a good fit to the observed decay (black, dashed lines in Fig. 3(d) and (e)). In all cases, the offset was positive suggesting that the dynamics are indeed not ergodic.

**Figure 4.**
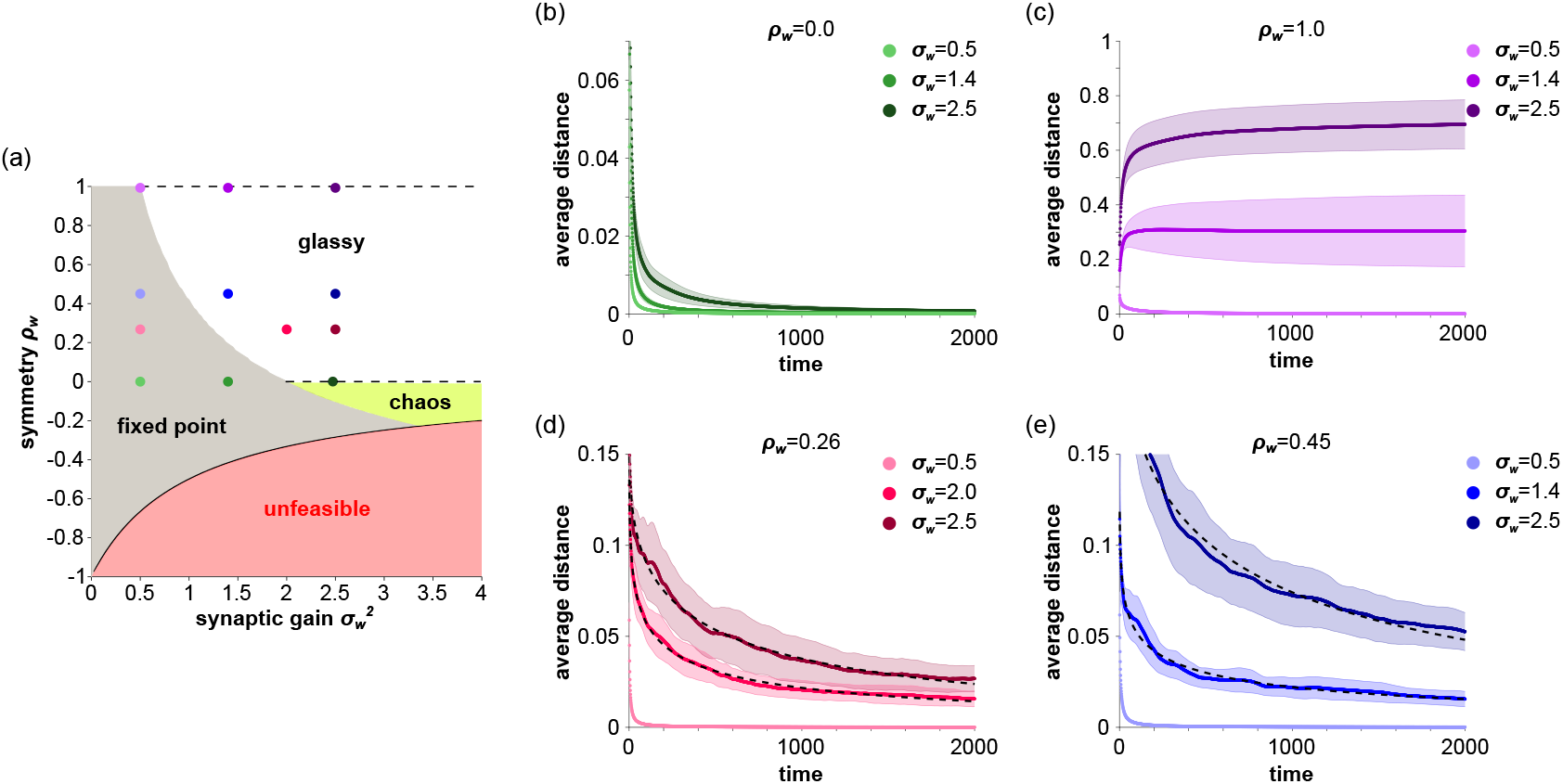
(a) Phase diagram showing the different dynamical regimes as a function of the synaptic gain 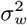 and the level of symmetry *ρ_w_*. Colored circles: representative points in the 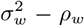 plane further illustrated in panels (b)-(e). The region marked *unfeasible* corresponds to pairs of 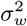 and *ρ_w_* that cannot be realized due to the positivity constraint for the synaptic efficacies, i.e., *w_ij_* > 0. (b)-(e) *D^ab^*, averaged over all pairs of initial conditions, as a function of time for different values of 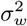 and *ρ_w_*. Shaded areas represent ±1 standard deviation from the mean. Colors identify the corresponding points in (a). Dashed black lines in (d) and (e) are least-squares fits to a stretched exponential (see main text for details).

In summary, we have shown that balanced networks of rate-based neurons exhibit three distinct phases. For low synaptic gain, there is a unique, stable fixed point regardless of the level of symmetry. The dynamics are chaotic and ergodic for random or anti-symmetric networks, while they are glassy for symmetric networks. In the glassy phase, settling in a fixed point becomes a very slow process and, accordingly, single neurons’ activity keeps fluctuating in time. From this point of view, the glassy phase resembles the chaotic one. The resemblance, however, is only superficial. Unlike in the chaotic phase, the memory of the initial conditions never fades away in the glassy phase.

Is there any evidence that the cortex operates in a glassy phase? Neurons in the cortex feature significant autocorrelation in spike counts separated by lags of seconds (see, e.g., [30]). This is a generic property of the glassy phase, which only requires positive levels of symmetry and large synaptic gains, as observed in the cortex. Alternative accounts [11, 12, 31–34] require significant fine-tuning of some parameter(s) (see [35] for a detailed discussion of this point). More concretely, the glassiness of the cortical dynamics could be directly assessed by looking at multi-point correlation functions [28, 29]. This appears feasible, given the availability of long, large-scale recordings of neuronal activity at the single-cell resolution.

Is there any computational advantage in operating in the glassy phase? It seems to us that there is no obvious disadvantage. Whatever benefit there is in operating in the chaotic phase, it can be only amplified by operating in the glassy phase, if only because of the broad spectrum of time scales and its robustness to changes in the synaptic gain and/or the level of symmetry. More interestingly, recent studies have shown that glassy materials can reliably perform useful computations, e.g., storing and retrieving memories, when driven cyclically [36]. These results, we believe, offer a fresh perspective for looking at *old* data, e.g., oscillations and short-term memory, possibly leading to new theoretical and experimental endeavors.

To conclude, we have shown that glassy dynamics can robustly occur in model systems whose dynamics are not described by an energy function. The underlying mechanism appears to be the same as the one leading to glassy dynamics in Hamiltonian systems, that is, the sudden appearance of a rugged *dynamical* (rather than energy) landscape as some control parameter is varied [19, 28, 29]. We note that our results support earlier proposals that, to have structurally-robust glassy behavior, it is sufficient for the system to have an exponential number of fixed points with a *broad* distribution of spectral gaps [37]. This suggests that glassy behavior could, indeed, be quite common in biology. If so, the theoretical and experimental tools developed in physics to understand glasses could provide a suitable paradigm, not just a useful metaphor, to understand quantitatively complex behavior in biological systems.

We are grateful to B. Cessac, V. Hakim, G. La Camera, Y. Loewenstein, J.-P. Nadal, M. Tsodyks and C. van Vreeswijk for many stimulating discussions, and to V. Hakim and G. La Camera for a crtitical reading of the manuscript. G.M. acknowledges the hospitality and the financial support of the Simons Center for Systems Biology, at the School of Natural Sciences, Institute for Advanced Study, Princeton, where this work has been finalized. This work was supported by grants ANR-19-CE16-0024-01 and ANR-20-CE16-0011-02 from the French National Research Agency and by a grant from the Simons Foundation (891851, G.M.). K.B. acknowledges a fellowship from the ENS Paris-Saclay and support from the Office of Naval Research N00014-17-1-2041 (to X.-J. Wang).

## Notes

### Competing Interest Statement

The authors have declared no competing interest.

