## Supplementary Information for "Glassy phase in dynamically-balanced neuronal networks"

### Supplemental Material:
### Glassy phase in dynamically balanced neuronal networks

Kevin Berlemont[1] and Gianluigi Mongillo[2,3]

[1]*Center for Neural Science, New York University, New York, NY 10003, USA*

[2]*Sorbonne Université, INSERM, CNRS,*
*Institut de la Vision, F-75012 Paris, France*

[3]*Centre National de la Recherche Scientifique (CNRS), Paris, France*

#### CONTENTS


I. Linear response 3

II. Mean-field theory 4

A. Self-consistency equations 7

III. Stability of the mean-field solution 8

A. Critical line 10

IV. Uniqueness of the *microscopic* fixed point associated to the mean-field solution 11

#### I. LINEAR RESPONSE

In this section, we derive the equations that determine the static susceptibility matrix, $\boldsymbol{\chi}$. The susceptibility matrix will be needed for the derivation of the mean-field theory in the next section. In a fixed point, the rates of the neurons satisfy the following system of equations

$$\nu_i = \phi(h_i) = \phi\left(\sqrt{N} - \frac{1}{\sqrt{N}}\sum_{j=1}^{N} w_{ij}\nu_j\right). \tag{1}$$

We now perturb the network by adding an external input, $\eta_k$, to each neuron $k$, and compute $\chi_{ik}$ as

$$\chi_{ik} = \frac{d\nu_i}{d\eta_k} = \frac{d\nu_i}{dh_i}\cdot\frac{dh_i}{d\eta_k} = \phi'_i\cdot\left(-\frac{1}{\sqrt{N}}\sum_{j=1}^{N} w_{ij}\chi_{jk} + \delta_{ik}\right), \tag{2}$$

where $\phi'_i$ is a short-hand notation for $\phi'(h_i)$, and $\delta_{ik}$ is the Kronecker symbol, i.e., $\delta_{ik} = 1$ for $i = k$ and $\delta_{ik} = 0$ for $i \neq k$. If all the $\eta_k$ are sufficiently small, and the fixed point is stable, the change in the rate of the neuron $i$ resulting from the perturbation is given by

$$\delta\nu_i = \sum_{k=1}^{N}\chi_{ik}\eta_k. \tag{3}$$

In a stable fixed point, we expect $\chi_{ii} = O(1)$ and $\chi_{ik} = O(1/\sqrt{N})$ for $i \neq k$ (see Eq. (2)). Thus, we write

$$\chi_{ik} = \delta_{ik}\chi_{ii}^{(0)} + \frac{\chi_{ik}^{(1)}}{\sqrt{N}} + \frac{\chi_{ik}^{(2)}}{N}, \tag{4}$$

and plug this into Eq. (2). For $i \neq k$, we obtain

$$\chi_{ik}^{(1)} = \phi'_i\cdot\left(-w_{ik}\chi_{kk}^{(0)} - \frac{1}{\sqrt{N}}\sum_{j=1}^{N} w_{ij}\chi_{jk}^{(1)} - \frac{1}{N}\sum_{j=1}^{N} w_{ij}\chi_{jk}^{(2)}\right). \tag{5}$$

In the above equation, the term on the l.h.s. as well as the first and third term on the r.h.s. are $O(1)$. The second term on the r.h.s., however, is $O(\sqrt{N})$ unless

$$\frac{1}{N}\sum_{j=1}^{N}\chi_{jk}^{(1)} = 0, \tag{6}$$

in which case it also becomes $O(1)$. Next, we multiply both sides of Eq. (5) by $1/N$ and sum over $i$. Assuming that Eq. (6) is satisfied, we obtain

$$\frac{1}{N}\sum_{i=1}^{N}\chi_{ik}^{(1)} = \langle\phi'\rangle\langle w\rangle\cdot\left(-\chi_{kk}^{(0)} - \frac{1}{N}\sum_{j=1}^{N}\chi_{jk}^{(2)}\right) = 0, \tag{7}$$

which implies

$$\frac{1}{N}\sum_{j=1}^{N}\chi_{jk}^{(2)} = -\chi_{kk}^{(0)}. \tag{8}$$

It is easy to see, using Eq. (4), Eq. (6) and Eq. (8), that in the limit $N\to+\infty$ the static susceptibilities satisfy the following *balance* condition

$$\sum_{i=1}^{N}\chi_{ik} = 0 \tag{9}$$

for each $k$.

We conclude this section by noting that the linear response of dynamically balanced networks presents some *exotic* feature. Consider, for instance, the effect of a homogeneous perturbation, i.e., $\eta_k = \eta$ for all $k$. Clearly, $\delta\nu_i \sim \eta$ for each $i$ (see Eq. (3)). However, the change in the network-averaged rate will be zero. In fact,

$$\frac{1}{N}\sum_{i=1}^{N}\delta\nu_i = \frac{1}{N}\sum_{i=1}^{N}\left(\eta\sum_{k=1}^{N}\chi_{ik}\right) = \frac{\eta}{N}\sum_{k=1}^{N}\left(\sum_{i=1}^{N}\chi_{ik}\right) = 0. \tag{10}$$

This is in sharp contrast with what would happen in non-balanced networks. There, the change in the network-averaged rate will also be proprtional to $\eta$, similarly to the changes in the individual rates. The precise cancellation of the linear responses to external perturbations, homogeneous or heterogeneous, is a dynamical consequence of the fact that, in a balanced network, the average rate cannot be changed by $O(1)$ changes in the external input.

#### II. MEAN-FIELD THEORY

In this section, we derive the mean-field theory for the model network described in the main text. In a fixed point, the input to neuron $i$, $h_i$, is given by

$$h_i = \sqrt{N} - \frac{1}{\sqrt{N}} \sum_{j=1}^{N} w_{ij}\nu_j, \tag{11}$$

with $\langle w \rangle = 1$ and

$$\nu_j = \phi(h_j) \equiv h_j \Theta(h_j), \tag{12}$$

where $\Theta(\cdot)$ is the Heaviside function. As explained in the main text, one cannot use the central limit theorem to determine the distribution of the $h_i$ in the fixed point (i.e., to evaluate the sum over $j$ in Eq. (11)) when there are pairwise correlations in the synaptic efficacies. This is because $w_{ij}$ and $\nu_j$ are not independent: $h_j$, hence $\nu_j$, depends on $w_{ji}$, and $w_{ji}$ and $w_{ij}$ are correlated. In fact, as a result of these correlations, the distribution of the $h_i$ in the fixed point is not Gaussian. Nevertheless, the distribution of the $h_i$ can be evaluated with the following *cavity* argument.

Let us rewrite the input to neuron $i$ as follows

$$h_i = \sqrt{N} - \frac{1}{\sqrt{N}} \sum_{j=1}^{N} w_{ij}(\nu_j^- + \delta\nu_j) \equiv u_i - \frac{1}{\sqrt{N}} \sum_{j=1}^{N} w_{ij}\delta\nu_j, \tag{13}$$

where $\nu_j^-$ is the activity of neuron $j$ (in the same network) when all outgoing connection from neuron $i$ have been removed, i.e., $w_{ji} = 0$ for all $j$; $\delta\nu_j$ is the change in activity of neuron $j$ due to the presence of neuron $i$; and we have defined $u_i$ and will refer to it as the *cavity* input in the following. The $w_{ij}$ and the $\nu_j^-$ are now independent, by construction, and we can use the central limit theorem to determine the distribution of the $u_i$ in the fixed point. Thus, for $N \to +\infty$,

$$u_i = \mu + z_i\sigma, \tag{14}$$

where $z_i$ is Gaussian with zero mean and unitary variance, and

$$\mu = \sqrt{N}\left(1 - \langle \nu^- \rangle\right), \tag{15}$$

$$\sigma^2 = \sigma_w^2 \langle (\nu^-)^2 \rangle. \tag{16}$$

Next, we deal with the *reaction term*, i.e., the last term on the r.h.s. of Eq. (13). The presence of neuron $i$ induces a small perturbation, of $O(1/\sqrt{N})$, in the input to neuron $j$. If the fixed point is stable, we can use linear response theory and write

$$\delta\nu_j = \sum_{k=1}^{N} \chi_{jk} \left( -\frac{1}{\sqrt{N}} w_{ki} \nu_i \right), \tag{17}$$

where $\chi_{jk}$ is the static susceptibility of the neuron $j$ to a change in the *input* to neuron $k$ (see Sec. I). Then, using Eq. (17),

$$\begin{aligned} -\frac{1}{\sqrt{N}} \sum_j w_{ij} \delta\nu_j &= \frac{\nu_i}{N} \sum_{j,k} w_{ij} \chi_{jk} w_{ki} \\ &= \nu_i \cdot \left( \frac{1}{N} \sum_{j=1}^{N} w_{ij} \chi_{jj} w_{ji} + \frac{1}{N} \sum_{(jk)} w_{ij} \chi_{jk} w_{ki} \right), \end{aligned} \tag{18}$$

where the sum in the last term on the r.h.s. is over all pairs $(j,k)$ with $j \neq k$. Note that the synaptic efficacies to/from neuron $i$ (e.g., $w_{ij}$ and $w_{ki}$) and the susceptibilities are uncorrelated. Then, the first sum in Eq. (18) is straightforward to evaluate, i.e.,

$$\frac{1}{N} \sum_{j=1}^{N} w_{ij} \chi_{jj} w_{ji} = \left(1 + \rho_w \sigma_w^2\right) \chi + O\left(\frac{1}{\sqrt{N}}\right), \tag{19}$$

where we have defined

$$\chi \equiv \frac{1}{N} \sum_{j=1}^{N} \chi_{jj}. \tag{20}$$

The second sum in Eq. (18) requires a little more care. We obtain

$$\frac{1}{N} \sum_{(jk)} w_{ij} \chi_{jk} w_{ki} = \frac{1}{N} \sum_{k=1}^{N} w_{ki} \left( \sum_{\substack{j=1 \\ j\neq k}}^{N} \chi_{jk} + \sum_{\substack{j=1 \\ j\neq k}}^{N} \delta w_{ij} \chi_{jk} \right) \tag{21}$$

$$= \frac{1}{N} \sum_{k=1}^{N} w_{ki} \left(-\chi_{kk} + z_{ik}\right) \tag{22}$$

$$= -\chi + O\left(\frac{1}{\sqrt{N}}\right). \tag{23}$$

In going from Eq. (21) to Eq. (22), for the first sum in the brackets, we have used Eq. (9), i.e.,

$$\sum_{j=1}^{N} \chi_{jk} = \chi_{kk} + \sum_{\substack{j=1 \\ j\neq k}}^{N} \chi_{jk} = 0 \implies \sum_{\substack{j=1 \\ j\neq k}}^{N} \chi_{jk} = -\chi_{kk}, \tag{24}$$

while, for the second sum in the brackets, we have used the facts that (i) the $\delta w_{ij}$ and the $\chi_{jk}$ are uncorrelated, and (ii) the $\chi_{jk}$ are $O(1/\sqrt{N})$ for $j \neq k$. Thus, if the variance of the $\chi_{jk}$ is finite (that is, the fixed point is stable), the second sum, $z_{ik}$, is $O(1)$ and normally distributed with mean 0. Eq. (23) then follows from Eq. (22) by noticing that the $z_{ik}$ are independent from the $w_{ki}$.

Putting all together, for the reaction term we obtain (in the limit $N \to +\infty$)

$$-\frac{1}{\sqrt{N}} \sum_{j} w_{ij} \delta\nu_j = \rho_w \sigma_w^2 \chi \nu_i \equiv R\nu_i, \tag{25}$$

and then, by requiring self-consistency for neuron $i$, we can determine $\nu_i$ as a function of $u_i$, i.e.,

$$\nu_i = \phi(u_i + R\nu_i) \implies \nu_i = \frac{u_i \Theta(u_i)}{1-R} \equiv \hat{\phi}(u_i). \tag{26}$$

##### A. Self-consistency equations

The statistics of the activity in the fixed point is completely determined by the mean and the variance of the cavity input, $\mu$ and $\sigma^2$ respectively, and by the average local susceptibility $\chi$. These can be determined by requiring self-consistency. In the limit $N \to +\infty$, the average activity in the network is determined by the balance condition, i.e., $\langle \nu \rangle = 1/\langle w \rangle = 1$. Thus

$$1 = \int Dz\, \hat{\phi}(\mu + z\sigma) = \frac{\sigma}{1-R} \left[G(x) - xH(x)\right] \tag{27}$$

where $Dz$ is the standard Gaussian measure, and we have defined $x = -\mu/\sigma$, and

$$G(x) \equiv \frac{1}{\sqrt{2\pi}} \exp\left(-\frac{x^2}{2}\right); \quad H(x) \equiv \int_x^{+\infty} dz G(z) = \int_x^{+\infty} Dz. \tag{28}$$

Similarly,

$$\sigma^2 = \sigma_w^2 \int Dz \left[\hat{\phi}(\mu + z\sigma)\right]^2 = \sigma_w^2 \left(\frac{\sigma}{1-R}\right)^2 \left[-xG(x) + (1+x^2)H(x)\right], \tag{29}$$

that is,

$$1 = \left(\frac{\sigma_w}{1-R}\right)^2 \left[-xG(x) + (1+x^2)H(x)\right]. \tag{30}$$

In writing Eq. (27) and Eq. (29), we have used the fact that $\langle (\nu^-)^n \rangle = \langle \nu^n \rangle$ for $N \to +\infty$. Finally, we need an equation for $\chi$. Using Eq. (26), we obtain

$$\chi_{ii} = \frac{d\nu_i}{du_i} = \frac{\Theta(u_i)}{1-R}, \tag{31}$$

and then, by definition,

$$\chi = \int Dz \, \frac{\Theta(\mu + z\sigma)}{1-R} = \frac{H(x)}{1-R}. \tag{32}$$

The self-consistency conditions Eq. (27), Eq. (30) and Eq. (32) can be further reduced to a single equation for $x$, in the following way. From Eq. (30),

$$1 - R = \sigma_w \left[-xG(x) + (1+x^2)H(x)\right]^{1/2} \equiv \sigma_w A(x) \tag{33}$$

and, using this into Eq. (32) (recall that $R = \rho_w \sigma_w^2 \chi$),

$$\sigma_w \left[A(x) + \rho_w \frac{H(x)}{A(x)}\right] = 1. \tag{34}$$

Eq. (34) has always only one solution. Once one knows the corresponding $x$, one computes $R$ from Eq. (33), and then $\sigma$ from Eq. (27). Finally, $\mu = -\sigma \cdot x$.

#### III. STABILITY OF THE MEAN-FIELD SOLUTION

The mean-field theory developed in Sec. II is based on the assumption that the fixed point is asymptotically stable. We can check the validity of this assumption by computing self-consistently the adjustments of the network activity due to the presence of neuron $i$. If these corrections are not defined or *nonphysical*, the fixed point is unstable.

Using the effective transduction function, $\hat{\phi}(\cdot)$ (see Eq. 26)), we can write the change in the activity of neuron $j$ due to the presence of neuron $i$, $\delta\tilde{\nu}_j$, as

$$\delta\tilde{\nu}_j = \hat{\phi}(u_j) - \hat{\phi}(u_j^-) = \hat{\phi}'(u_j)\delta\tilde{u}_j = \hat{\phi}'(u_j)\left(-\frac{1}{\sqrt{N}}\sum_k w_{jk}\delta\tilde{\nu}_k - \frac{1}{\sqrt{N}}w_{ji}\nu_i\right), \tag{35}$$

where $u_j$ and $u_j^-$ are, respectively, the cavity input to neuron $j$ in the network with, and without, the neuron $i$. The $w_{jk}$ and the $\delta\tilde{\nu}_k$ are, by construction, uncorrelated. Then, the corrections to the cavity input are normally distributed with mean, $\langle\delta\tilde{u}\rangle$, and variance, $\Delta^2$, given by

$$\langle\delta\tilde{u}\rangle = -\frac{1}{\sqrt{N}}\left(N\langle\delta\tilde{\nu}\rangle + \nu_i\right), \tag{36}$$

$$\Delta^2 = \sigma_w^2\left(\langle\delta\tilde{\nu}^2\rangle + \frac{\nu_i^2}{N}\right). \tag{37}$$

The cavity input to neuron $j$ and its correction, $\delta\tilde{u}_j$, are not independent because they are determined by the same set of incoming synaptic efficacies, $w_{jk}$. Their covariance, $\xi$, is given by

$$\xi = \langle\left(u_j - \mu\right)\left(\delta\tilde{u}_j - \langle\delta\tilde{u}\rangle\right)\rangle = \sigma_w^2\langle\nu\delta\tilde{\nu}\rangle. \tag{38}$$

In a stable fixed point, we expect $\xi = O\left(1/\sqrt{N}\right)$. Thus, we write

$$u_j = \mu + x_j\sigma + z_j\sqrt{\xi} \quad \text{(to leading order)}, \tag{39}$$

$$\delta\tilde{u}_j = \langle\delta\tilde{u}\rangle + y_j\sqrt{\Delta^2 - \xi} + z_j\sqrt{\xi}, \tag{40}$$

where $x_j$, $y_j$ and $z_j$ are independent Gaussian variables with mean 0 and variance 1. Using Eqs. (39)-(40) we can compute $\langle\delta\tilde{\nu}\rangle$ via Eq. (35). We obtain

$$\langle\delta\tilde{\nu}\rangle = \langle\hat{\phi}'(u)\delta\tilde{u}\rangle = \langle\hat{\phi}'\rangle\langle\delta\tilde{u}\rangle + \langle\hat{\phi}''\rangle\xi = -\frac{1}{\sqrt{N}}\chi\left(N\langle\delta\tilde{\nu}\rangle + \nu_i\right), \tag{41}$$

where we have used $\langle\hat{\phi}'\rangle = \chi$ and $\langle\hat{\phi}''\rangle = 0$ (see Eqs. (31)-(32)). Similarly, for $\langle\nu\delta\tilde{\nu}\rangle = \langle\hat{\phi}(u)\hat{\phi}'(u)\delta\tilde{u}\rangle$ we obtain

$$\begin{aligned}\langle\nu\delta\tilde{\nu}\rangle &= \langle\hat{\phi}\hat{\phi}'\rangle\langle\delta\tilde{u}\rangle + \left(\langle\hat{\phi}'^2\rangle + \langle\hat{\phi}\hat{\phi}''\rangle\right)\xi \\ &= -\frac{1}{\sqrt{N}}\langle\hat{\phi}\hat{\phi}'\rangle\left(N\langle\delta\tilde{\nu}\rangle + \nu_i\right) + \sigma_w^2\chi_{nl}\langle\nu\delta\tilde{\nu}\rangle,\end{aligned} \tag{42}$$

where have used $\langle\hat{\phi}\hat{\phi}''\rangle = 0$ and have defined $\chi_{nl} = \langle\hat{\phi}'^2\rangle$. Eqs. (41)-(42) are a set of self-consistent (linear) equations that determine $\langle\delta\tilde{\nu}\rangle$ and $\langle\nu\delta\tilde{\nu}\rangle$ as a function of the *uniform* component of the perturbation, i.e., -$\nu_i/\sqrt{N}$. They are easily solved by posing

$$\delta\tilde{\nu}_j = \frac{\delta\nu_j}{\sqrt{N}} + \frac{\delta\nu_j^{(1)}}{N}. \tag{43}$$

We find $\langle\delta\nu\rangle = 0$ and

$$\begin{cases} -\chi\left(\langle\delta\nu^{(1)}\rangle + \nu_i\right) = 0 \\ -\langle\hat{\phi}\hat{\phi}'\rangle\left(\langle\delta\nu^{(1)}\rangle + \nu_i\right) + \left(-1 + \sigma_w^2\chi_{nl}\right)\langle\nu\delta\nu\rangle = 0 \end{cases} \tag{44}$$

whose unique solution is $\langle\delta\nu^{(1)}\rangle = -\nu_i$ and $\langle\nu\delta\nu\rangle = 0$, provided

$$1 - \sigma_w^2\chi_{nl} \neq 0. \tag{45}$$

Using $\langle\delta\nu\rangle = 0$ and $\langle\delta\nu^{(1)}\rangle = -\nu_i$ in Eq. (35) we obtain

$$\delta\nu_j = -\hat{\phi}'(u_j)\left(\delta w_{ji}\nu_i + \frac{1}{\sqrt{N}}\sum_k \delta w_{jk}\delta\nu_k\right). \tag{46}$$

Taking the square and averaging over $j$, we finally obtain

$$\langle\delta\nu^2\rangle = \frac{\sigma_w^2\chi_{nl}}{1 - \sigma_w^2\chi_{nl}}\nu_i^2 \equiv R_{nl}\nu_i^2. \tag{47}$$

Clearly, in a *physical* solution $\langle\delta\nu^2\rangle$ must be positive and finite, which requires

$$\sigma_w^2\chi_{nl} < 1. \tag{48}$$

Obviously, if Eq. (48) is satisfied, condition Eq. (45) is also satisfied.

##### A. Critical line

The mean-field solution, that is the solution of Eq. (34), is stable when Eq. (48) is satisfied. Recalling Eq. (31) and the definition of $\chi_{nl}$, Eq. (48) is satisfied when

$$\frac{\sigma_w^2 H(x)}{(1-R)^2} = \frac{H(x)}{A^2(x)} < 1, \tag{49}$$

because in the mean-field solution $(1-R)^2 = \sigma_w^2 A^2(x)$ (see Eq. (33)). Eq. (49) implies that $x$ is constant on the critical line, i.e., $x^{(c)} = 0$ along the critical line. Using this information into Eq. (34), we obtain the equation for the critical line

$$\sigma_w^{(c)} \left(1 + \rho_w^{(c)}\right) = \sqrt{2}. \tag{50}$$

Note that the susceptibility $\chi$ (and hence the reaction term $R$) is finite on the critical line. This is related to the fact that there is no change in the average activity level when crossing the critical line. The instability, instead, is signaled by the divergence of the non-linear reaction term $R_{nl}$, defined in Eq. (47).

#### IV. UNIQUENESS OF THE *MICROSCOPIC* FIXED POINT ASSOCIATED TO THE MEAN-FIELD SOLUTION

The mean-field theory determines the marginal distribution of the synaptic inputs $h_i$ in a fixed point of the network dynamics. This is, by its very nature, a *macroscopic* description of the network activity (in a fixed point). By contrast, a *microscopic* description consists of the $N$-vector of the $h_i$ in a fixed point. Any permutation of this vector, which corresponds in fact to a *different* solution of Eq. (11), would have the same marginal statistics and, hence, would be consistent with the mean-field solution. In other words, multiple microscopic states can be associated to the mean-field solution.

Let us suppose that there exists $n$ different fixed points, i.e., solutions of Eq. (11). Then, the distribution of the cavity inputs to a *given* neuron in the *different* fixed points is described by an $n$-variate Gaussian in the mean-field approximation. Mean(s) and variance(s) are known from the mean-field solution. They are the same in all fixed points because the mean-field solution is unique. We need to compute the off-diagonal elements of the co-variance matrix, $\langle \delta u^a \delta u^b \rangle$. These are given by

$$\langle \delta u^a \delta u^b \rangle \equiv \frac{1}{N}\sum_{i=1}^{N} \delta u_i^a \delta u_i^b = \frac{1}{N}\sum_{i=1}^{N}\left[\left(\frac{1}{\sqrt{N}}\sum_{j=1}^{N}\delta w_{ij}\nu_j^a\right)\cdot\left(\frac{1}{\sqrt{N}}\sum_{k=1}^{N}\delta w_{ik}\nu_k^b\right)\right] \tag{51}$$

$$= \frac{1}{N}\sum_{i=1}^{N}\left[\frac{1}{N}\sum_{j=1}^{N}\delta w_{ij}^2 \nu_j^a \nu_j^b + \frac{1}{N}\sum_{\substack{j,k=1\\ j\neq k}}^{N}\delta w_{ij}\delta w_{ik}\nu_j^a \nu_k^b\right] \tag{52}$$

$$= \sigma_w^2 \frac{1}{N}\sum_{j=1}^{N}\nu_j^a\nu_j^b + \frac{1}{N}\sum_{\substack{j,k=1\\ j\neq k}}^{N}\left(\frac{1}{N}\sum_{i=1}^{N}\delta w_{ij}\delta w_{ik}\right)\nu_j^a\nu_k^b \tag{53}$$

$$= \sigma_w^2 \langle \nu^a \nu^b \rangle + O\left(\frac{1}{\sqrt{N}}\right). \tag{54}$$

Thus, in the limit $N \to \infty$, we can write

$$\delta u_i^a = \sigma_w \cdot \left(x_i^a \sqrt{\langle \nu^2 \rangle - \langle \nu^a \nu^b \rangle} + z_i^{ab}\sqrt{\langle \nu^a \nu^b \rangle}\right), \tag{55}$$

$$\delta u_i^b = \sigma_w \cdot \left(x_i^b \sqrt{\langle \nu^2 \rangle - \langle \nu^a \nu^b \rangle} + z_i^{ab}\sqrt{\langle \nu^a \nu^b \rangle}\right), \tag{56}$$

where $x_i^a$, $x_i^b$ and $z_i^{ab}$ are independent Gaussian variables with zero mean and unitary variance. Finally, using the effective transduction function $\hat{\phi}(\cdot)$, we obtain a self-consistent equation that determines $\langle \nu^a \nu^b \rangle$, i.e.,

$$\langle \nu^a \nu^b \rangle = \iiint Dx^a Dx^b Dz^{ab} \left[\hat{\phi}\left(\mu + \sigma_w \cdot \left(x^a \sqrt{\langle \nu^2 \rangle - \langle \nu^a \nu^b \rangle} + z^{ab}\sqrt{\langle \nu^a \nu^b \rangle}\right)\right)\right. \\ \left. \times\, \hat{\phi}\left(\mu + \sigma_w \cdot \left(x^b \sqrt{\langle \nu^2 \rangle - \langle \nu^a \nu^b \rangle} + z^{ab}\sqrt{\langle \nu^a \nu^b \rangle}\right)\right)\right]. \tag{57}$$

Eq. (57) has always one solution, that is $\langle \nu^a \nu^b \rangle = \langle \nu^2 \rangle$, as it is easy to check. By solving numerically Eq. (57), we find that $\langle \nu^a \nu^b \rangle = \langle \nu^2 \rangle$ is, in fact, the only solution as long as the mean-field solution is stable. This means that, if there are multiple fixed points, they can only differ at a vanishingly small number of $h_i$, when $N \to \infty$. Thus, there is only one *microscopic* state (i.e., $n = 1$) associated to the mean-field solution.
